# Ferrosomes are iron storage organelles formed by broadly conserved gene clusters in bacteria and archaea

**DOI:** 10.1101/2020.01.10.902569

**Authors:** Carly R. Grant, Arash Komeili

## Abstract

Cellular iron homeostasis is vital and maintained through tight regulation of iron import, efflux, storage, and detoxification^1–3^. The most common modes of iron storage employ proteinaceous compartments that are composed of ferritin or related proteins^4,5^. While lipid-bounded iron compartments have also been described, the basis for their formation and function remains unknown. Here, we focus on one such compartment, the ferrosome, which had been previously observed in the anaerobic bacterium *Desulfovibrio magneticus*^6^. We identify three ferrosome-associated (Fez) proteins, encoded by a putative operon, that are associated with and responsible for forming ferrosomes in *D. magneticus*. Fez proteins include FezB, a P_1B-6_-ATPase found in phylogenetically and metabolically diverse species of bacteria and archaea with anaerobic lifestyles. In the majority of these species, two to ten genes define a cluster that encodes FezB. We show that two other species, *Rhodopseudomonas palustris* and *Shewanella putrefaciens*, make ferrosomes in anaerobic conditions through the action of their six-gene *fez* operon. Additionally, we find that the *S. putrefaciens fez* operon is sufficient for ferrosome formation in *Escherichia coli*. Using *S. putrefaciens* as a model, we find that ferrosomes likely play a role in the anaerobic adaptation to iron starvation. Overall, this work establishes ferrosomes as a new class of lipid-bounded iron storage organelles and sets the stage for studying ferrosome formation and structure in diverse microorganisms.

---

*D. magneticus* is an anaerobic sulfate-reducing bacterium and an emerging model organism for studying the natural diversity of magnetite biomineralization within an organelle termed the magnetosome^7,8^. Independent of magnetosomes, *D. magneticus* makes subcellular electron-dense granules rich in iron, phosphorus, and oxygen that are enclosed by a lipid-like membrane^6^. These granules, which we propose to name ‘ferrosomes’ for ‘iron bodies’, are visible by transmission electron microscopy (TEM) after *D. magneticus* transitions out of iron starvation with the supplementation of iron at a range of concentrations (Extended Data Fig. 1)^6^. We had previously found that the iron accumulated in ferrosomes is not sufficient for magnetosome formation and that magnetosome genes are not required for ferrosome formation^6,9^. While these studies support the hypothesis that the ferrosome is a distinct organelle in *D. magneticus*, the molecular basis for ferrosome formation and function has remained a mystery.

To understand the mechanistic basis of ferrosome formation, we isolated ferrosomes from cell lysates through a sucrose cushion and used mass spectrometry to identify their associated proteins (Extended Data Fig. 2a-c). Relative protein quantification revealed three proteins highly enriched in the ferrosome fraction, DMR_28330 (FezB), DMR_28340 (FezC), and DMR_28320 (FezA), that are encoded by genes arranged in a putative operon, *fezABC* (Fig. 1a, b) (gene prefix given for the phonetic pronunciation ‘ferrozome’). Of these three proteins, only FezB has a functional annotation: a heavy metal-transporting P_1B_-ATPase. P_1B_-ATPases are a large family of integral membrane proteins that transport metals across membranes using the energy of ATP hydrolysis^10^. FezB falls within the P_1B-6_-ATPase group, an uncharacterized subfamily with unique transmembrane topology and a possible role in iron transport based on genomic context in several species^11^. FezB has the cytoplasmic domains characteristic of all P_1B_-ATPases and unique motifs in the transmembrane domains responsible for metal specificity^10,11^ (Fig. 1c, Extended Data Fig. 3). FezC has an N-terminal heavy-metal associated (HMA) domain annotation and two predicted transmembrane domains while FezA has a hydrophobic N-terminal region (Extended Data Fig. 2d). The putative transmembrane domains of FezA, FezB, and FezC are consistent with our previous observations that ferrosomes are surrounded by a lipid-like membrane^6^. Additionally, the characteristics of metal binding and transport domains imply that the *fez* operon may be the geneticblueprint for ferrosome formation and function.

**Fig. 1.**
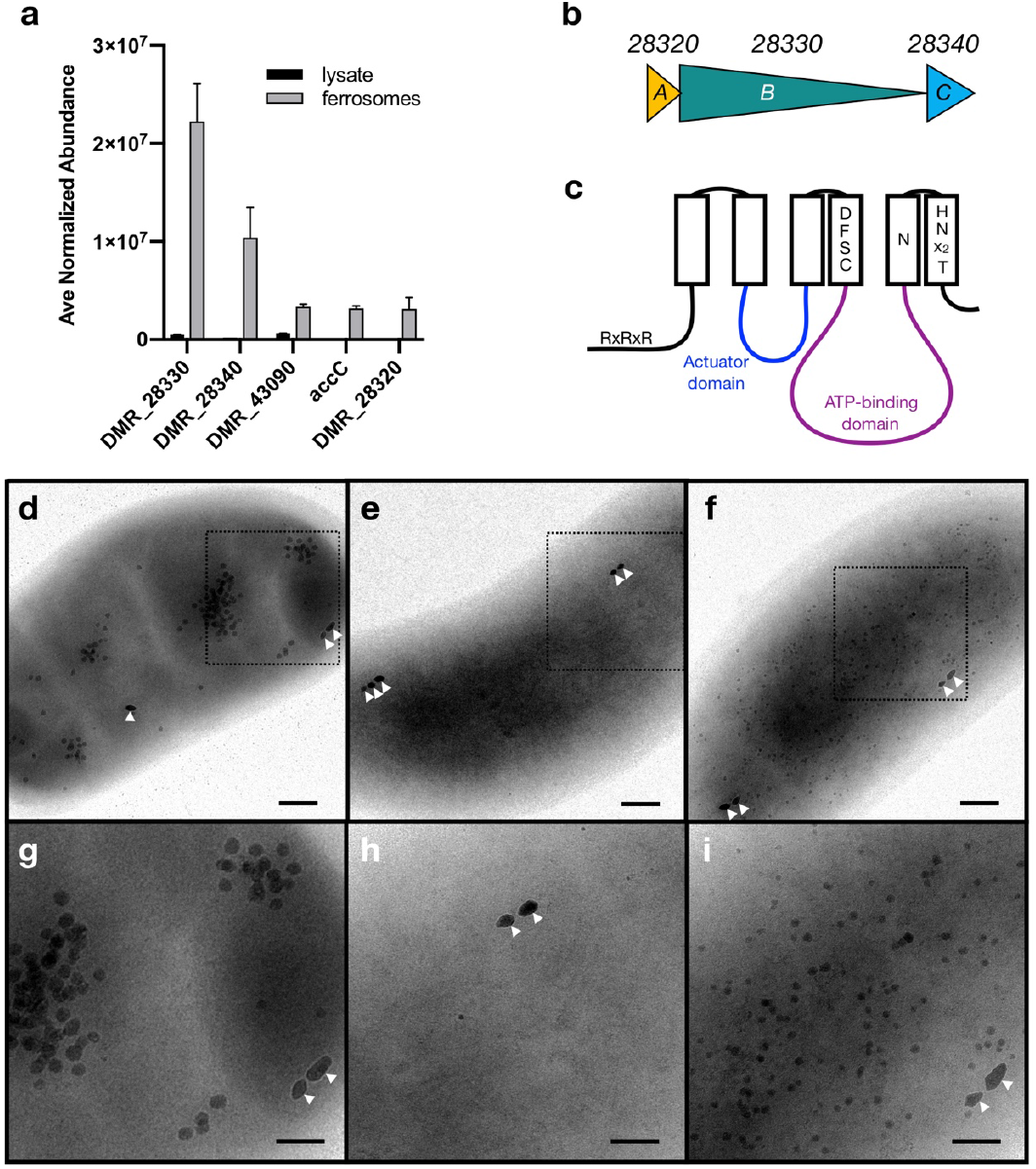
Proteins enriched with ferrosomes isolated from *D. magneticus* are essential for ferrosome formation. (a) Average normalized abundance of five proteins most highly enriched with isolated ferrosomes compared with the whole cell lysate, as detected by LC-ESI/MS. (b) DMR_28320-40 are encoded by genes that are arranged in a putative operon, *fezABC*. (c) Schematic of FezB. FezB has the conserved actuator and ATP-binding domains found in all P_1B_-ATPases and six putative trannsmembrane domains (rectangles). Signature motifs in the N-terminal domain and transmembrane domains 4-6, which may be involved in metal-binding and transport, are shown. Details of this schematic are based on the alignments shown in Extended Data Fig. 3. (d-i) Micrographs of *D. magneticus* strains after transitioning out of iron starvation. WT *D. magneticus* (d, g) has visible ferrosomes that are not found in the Δ*fezBC*_*Dm*_ strain (e, h). Complementation with *fezABC* expressed in *trans* rescues the phenotype (f, i). Micrographs in g-i are insets of d-f. White carets indicate magnetosomes. Scale bars, 200 nm; insets, 100 nm.

To test this hypothesis, we deleted the *D. magneticus fezB* and *fezC* genes through allelic replacement with a streptomycin-resistance cassette. The resulting mutant, Δ*fezBC_Dm_*, could still form magnetosomes but was unable to form ferrosomes—a phenotype that could be complemented *in trans* (Fig. 1d-i). In addition, expression of *fezABC*_*Dm*_ *in trans* in either the wild-type (WT) or in the Δ*fezBC*_*Dm*_ mutant led to constitutive ferrosome production in iron replete medium with no effect on magnetosome formation (Extended Data Fig. 4). Thus, ferrosomes are a structurally and genetically distinct organelle in *D. magneticus*.

We next asked if other organisms were also capable of forming ferrosomes. Phylogenetic analysis of FezB revealed a clear group of its homologs that share signature motifs in the metal binding transmembrane domains (D[Y/F]SCA and HNxxT, respectively) which define the P_1B-6_-ATPase subgroup^11^ (Fig. 2a, Supplementary Table 1). While FezB homologs lack a known cytoplasmic N-terminal metal binding domain, we found a notable “R-rich” motif containing two or more arginine residues spaced by a variable residue (e.g. RxR or RxRxR) in the N-terminus of the majority of FezB homologs and also related P_1B_-ATPases (Fig. 2a, Supplementary Table 1), including CtpC, a metal transporter that contributes to *M. tuberculosis* virulence^12,13^. FezB homologs are found in diverse species of bacteria and archaea that inhabit a range of environmental and host-associated habitats. While metabolically diverse, the majority of these species are strict or facultative anaerobes (Supplementary Table 2).

**Fig. 2.**
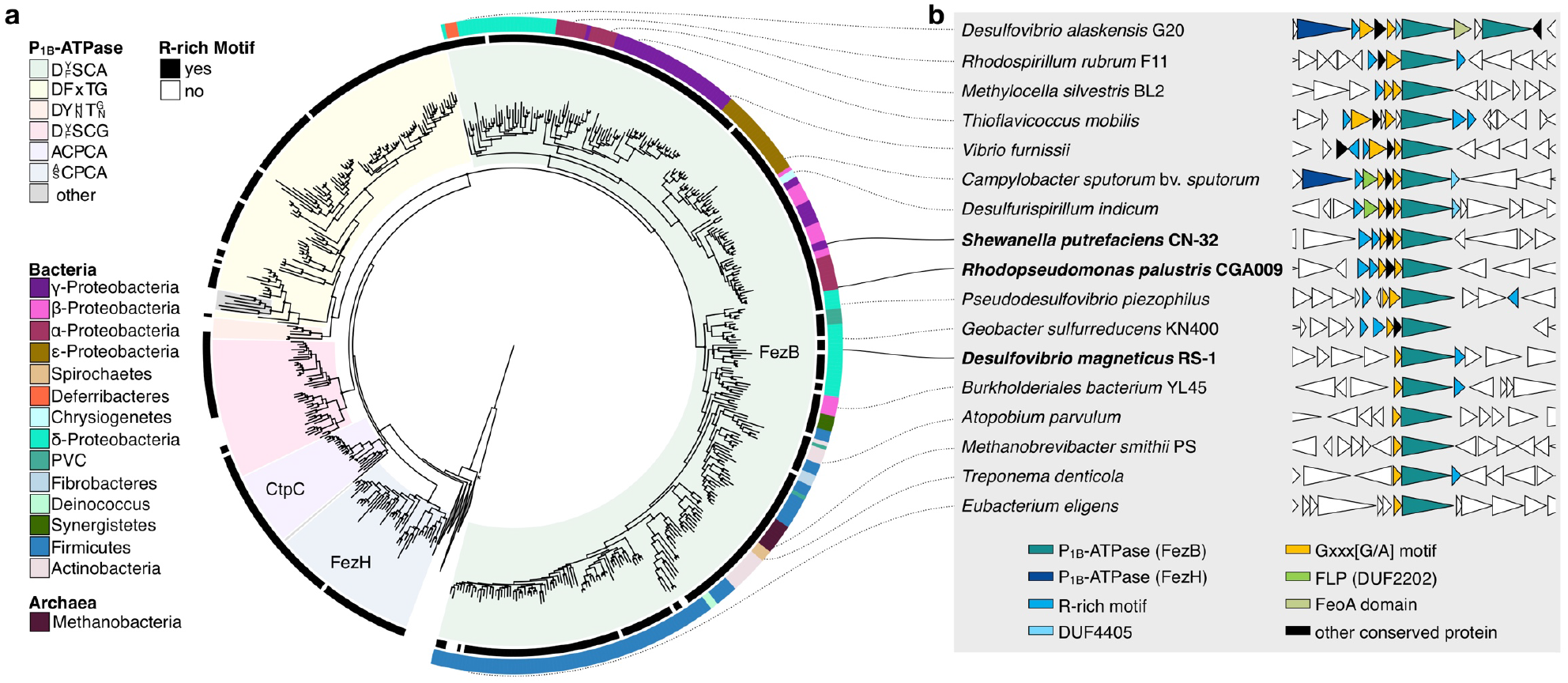
Phylogenetic analysis of FezB. (a) A maximum likelihood tree shows the relationship of FezB with other P_1B_ - ATPases. FezB has a signature motif in the metal-binding transmembrane domain that distinguishes it from other subgroups of P_1B_-ATPases (color ranges). The N-terminal R-rich motif is not specific to FezB, as related P_1B_-ATPases also have this motif (internal black color strip). The phylum or superphylum of organisms with a FezB homolog are indicated with the external color strip. The collapsed clades contain P_1B_-ATPases that lack an R-rich motif, including CopA, CopB, ZntA, and PfeT. Bootstraps >70% are indicated with black circles. (b) FezB is encoded by conserved gene clusters. Representative species with a FezB homolog and their *fez* gene cluster arrangement are shown. The lines connecting to the phylogenetic tree indicate the position of the FezB homolog for each species. The bacteria investigated in this study are in bold. Conserved *fez* gene cluster-encoded proteins are colored.

In most species, *fezB* lies in a conserved gene cluster (Extended Data Fig. 5a). Upon closer inspection, we found that nearly all *fez* gene clusters have one or more proteins that, like FezA, have a hydrophic region with a conserved GxxxG motif (Fig. 2b, Extended Data Fig. 5b, 6b, Supplementary Table 4). GxxxG motifs are common in transmembrane domains where they may facilitate protein-protein interactions and have even been shown to induce local curvature and tubulation of membranes^14–16^. Many *fez* gene clusters also encode one or more proteins with an N-terminal R-rich motif similar to that found in FezB (Extended Data Fig. 5b, 6a, Supplementary Table 5). These proteins comprise both soluble and membrane proteins, including FezC (Extended Data Fig. 6a). In some of the larger *fez* gene clusters, we discovered a second uncharacterized P_1B_-ATPase (FezH) with an R-rich motif and distinct transmembrane metal binding sites (Fig. 2, Extended Data Fig. 6a, Supplementary Table 3). Conserved proteins also include a DUF4405 protein with homology to the membrane domains of FezC, a FeoA-domain containing, and a DUF2202 ferritin-like protein with a C-terminal GxxxG motif (Fig. 2, Extended Data Fig. 5b, 6, Supplementary Table 6). The presence of these genes, as well as the genomic association of *fez* gene clusters with iron homeostasis genes^11^ (Extended Data Fig. 5c, Supplementary Table 7), supports a model in which a complex of Fez proteins transport iron into ferrosomes for storage.

The broad phylogenetic distribution of *fez* gene clusters suggests that diverse species of bacteria and archaea might be capable of forming ferrosomes. Since most of these organisms are uncultured or difficult to manipulate in the lab, we searched for culturable bacteria with established tools for genetic manipulation. *S. putrefaciens* is particularly conspicuous amongst these bacteria as it has been shown to form membrane-enclosed electron-dense granules consisting of mixed-valence iron, phosphorus, and oxygen^17,18^. These granules could not be found in several other *Shewanella* species^18^. Amongst the *Shewanella* species tested in these studies, *S. putrefaciens* is the only one with a putative *fez* operon (Fig. 2, Fig. 3a). Thus, we hypothesized that the iron-containing granules observed in previous studies are analogous to ferrosomes made by *D. magneticus*.

**Fig. 3.**
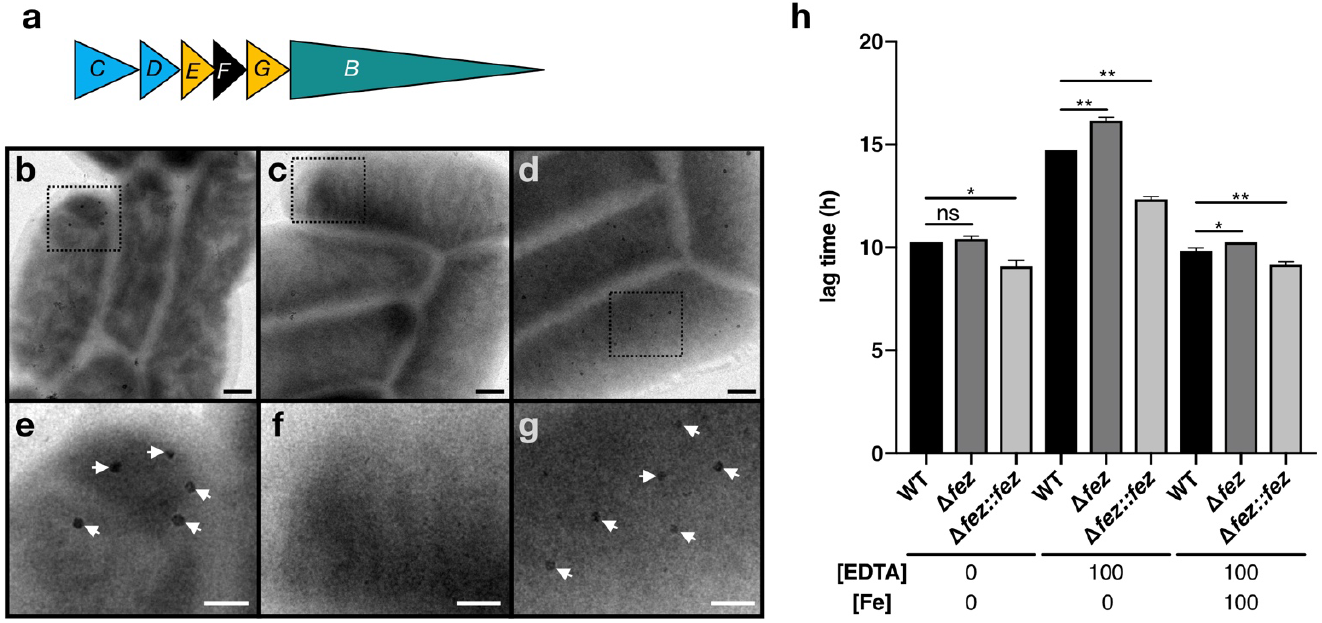
*fez* genes are essential for ferrosome formation and function in *S. putrefaciens*. (a) The *S. putrefaciens* six gene *fez* operon encodes FezB and FezC in addition to two GxxxG motif-containing proteins of unknown function, FezE and FezG, a soluble chaperone-like protein, FezD, and a putative protein with two transmembrane domains, FezF. (b-g) Micrographs of *S. putrefaciens* strains respiring HFO. WT *S. putrefaciens* makes ferrosomes (b, e) that are not found in the Δ*fez*_*Sp*_ strain (c, f). This phenotype is recued in the complementation strain, Δ*fez_Sp_∷fez*_*Sp*_ (d, g). Micrographs e-g are insets of b-d. White arrows indicate ferrosomes. Scale bars, 200 nm; insets, 100 nm. (h) Compared to WT, the *Δfez*_*Sp*_ mutant has a significantly longer lag time when grown in medium supplemented with EDTA (100 *μ*M) while the Δ*fez_Sp_∷fez*_*Sp*_ strain has a significantly shorter lag time. Adding back equimolar amounts of iron (100 *μ*M ferrous iron) rescues the phenotype. Data presented are averages of 3 independent cultures; error bars indicate the standard deviations. Statistical significance determined using Welch’s t-test. ns, not significant; *, p<0.05, **, p<0.01.

As described in previous work, we found that *S. putrefaciens* forms electron-dense granules when respiring hydrous ferric oxide (HFO) or fumarate in anaerobic growth medium supplemented with iron (Fig. 3b, e, Extended Data Fig. 7a, b)^17,18^. Likewise, *R. palustris* CGA009, which has a similar *fez* operon to *S. putrefaciens*, forms electron-dense granules resembling ferrosomes when grown photoheterotrophically in anaerobic medium supplemented with iron (Extended Data Fig. 8b). This is in accordance with previous proteomics and transcriptomic studies that show *fez* genes are expressed during anaerobic conditions in *R. palustris* strains CGA009 and TIE-1^19–21^. Notably, *S. putrefaciens* and *R. palustris* fail to produce ferrosomes when grown aerobically (Extended Data Fig. 8a)^17^. To confirm that the granules in *S. putrefaciens* and *R. palustris* are ferrosomes, we made markerless deletions of their *fez* gene clusters (Δ*fez*_*Sp*_ and Δ *fez*_*Rp*_, respectively). Mutants lacking the *fez* genes no longer made granules and complementation by expressing the *fez* genes *in trans* rescued the phenotype (Fig. 3c-g, Extended Data Fig. 7, 8).

We next asked whether or not *fez* genes can lead to ferrosome formation in a naïve host. To answer this question, we heterologously expressed the *S. putrefaciens fez* gene cluster on a plasmid in *E. coli*. When grown anaerobically in medium supplemented with iron, *E. coli* expressing *fez*_*Sp*_ (*E. coli / fez*_*Sp*_^+^) had a visibly dark pellet whereas the control cultures had a white pellet (Fig. 4a, b). TEM revealed electron-dense granules in *E. coli / fez*_*Sp*_^+^ cells that formed a dark pellet (Fig. 4). The granules have a diameter of around 20 nm which is nearly double that of the proteinaceous iron storage compartments found naturally in *E. coli*^1^. Taken together, these results show that *fez* genes are necessary and sufficient for ferrosome formation in diverse bacteria.

**Fig. 4.**
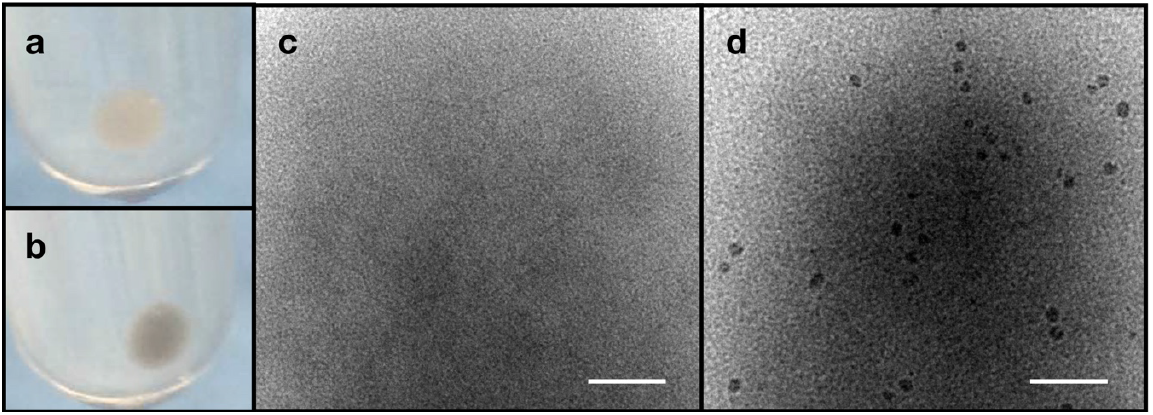
*E. coli* makes ferrosomes when expressing the *S. putrefaciens fez* genes heterologously. *E. coli* / *fez*_*Sp*_^+^ has a visibly dark cell pellet when grown anaerobically in growth medium supplemented with iron (b). (c, d) Transmission electron micrographs of *E. coli* strains grown anaerobically in growth medium supplemented with iron show electron-dense granules in *E. coli* / *fez*_*Sp*_^+^ (d). No granules are visible in *E. coli* harboring a control plasmid (c), which has a white cell pellet (a). Scale bars, 100 nm.

The genetic components of *fez* gene clusters and the patterns of ferrosome formation point to a role for this organelle in iron homeostasis. In other systems, iron storage compartments are important for surviving iron starvation. Using *S. putrefaciens* as a model, we found that addition of the iron chelator EDTA impaired aerobic and anaerobic growth for both the WT and the Δ*fez*_*Sp*_ strains. When grown aerobically, where no ferrosomes are formed^17^, the WT and Δ*fez*_*Sp*_ strains showed no difference in growth (Extended Data Fig. 9a). When grown anaerobically with EDTA, the WT and Δ*fez*_*Sp*_ mutant had a similar increase in doubling time, however the Δ*fez*_*Sp*_ mutant had a significantly longer lag phase compared to WT (Fig. 3h, Supplementary Table 8). This phenotype is specifically due to iron limitation, since it can be rescued by addition of equimolar concentrations of iron (Fig. 3h). In contrast, the complementation strain, Δ*fez*_*Sp*_∷*fez*_*Sp*_, which overproduces ferrosomes, had a significantly shorter lag time than the WT when grown anaerobically with or without EDTA (Fig. 3h, Supplementary Table 8). Overall, these results mirror the iron storage defect reported in the *E. coli* ferritin mutant during aerobic growth^22^. They are also consistent with recent findings that lag phase is a growth period dominated by accumulation of metals, such as iron, needed for the heavy enzymatic burden of exponential phase^23^. Therefore, we propose that ferrosomes likely function to store iron during anaerobic metabolism which can be accessed under severe iron starvation conditions.

In summary, our study reveals the genetic requirement for ferrosome formation and provides evidence that it functions as an iron storage organelle during anaerobic metabolism. Our findings that membrane proteins are associated with and required for ferrosome function support two independent studies that found lipid-like membranes surrounding ferrosomes^6,17^. This is in stark contrast to all other previously described bacterial and archaeal systems that depend on proteinaceous compartments for iron storage^4,5^. We propose that FezB has an unusual subcellular localization where it maintains the classic function of P_1B_-ATPases: to transport metals across membranes. Unlike most bacterial and archaeal P_1B_-ATPases that maintain metal homeostasis at the cell membrane, FezB transports iron into cytoplasmic ferrosome vesicles where it may be accessed when iron is limiting. This unique function may also apply to related P_1B_-ATPases that we identified in our phylogenetic analysis of FezB. While this study focused on environmental bacteria, iron storage may be a universal function of ferrosomes, including in host-associated bacteria. This hypothesis is supported by several unrelated studies in multiple bacteria that show *fez* gene expression is upregulated in low iron environments^24–29^, including during infection by *Clostridium difficile*^30^. In the future, ferrosomes may prove to be a novel drug target for combating pathogenic bacteria. They may also be platforms for synthetic biomining and bioremediation applications that leverage their metal-accumulating capabilities.

## Methods

### Strains, media, and, growth conditions

The bacterial strains used in this study are listed in Supplementary Table 9. All aerobic cultures were grown with continuous shaking at 250 rpm. Anaerobic cultures were grown at 30°C in an anaerobic glovebox or in sealed Balch tubes with a N_2_ headspace containing medium that was degassed with N_2_. Ferrous iron stocks were prepared by dissolving 1 M FeSO_4_ in 0.1 N HCl, which was subsequently stored in an anaerobic glovebox. Stocks of ferric malate were prepared as 20 mM FeCl_3_/60 mM malate. If needed, nitrilotriacetic acid (NTA) disodium salt was added to the ferrous iron to prevent precipitation of iron in the growth medium. NTA alone did not affect cellular growth.

*D. magneticus* strains were grown at 30°C anaerobically in RS-1 growth medium (RGM), as described previously^6,9^. For growth in iron replete medium, 100 *μ*M ferric malate was added to RGM prior to inoculation. For growth in iron limited medium (IL-RGM), iron was omitted from RGM and all glassware was soaked in oxalic acid for one to two days, as described previously^6^. To starve cells of iron, cultures were passaged in IL-RGM, as described previously^6^, or washed with IL-RGM prior to inoculation. To induce ferrosome formation, iron-starved cells were grown anaerobically in IL-RGM until they reached log-phase (OD_650_ ~0.1), at which point ferric malate was added to the cultures at a concentration of 100 *μ*M^6^.

*S. putrefaciens* strains were grown aerobically at 30°C in Luria-Bertani (LB) broth or anaerobically at 30°C in LB broth supplemented with 10 mM lactate and 10 mM fumarate or 40 mM hydrous ferric oxide (HFO). HFO was prepared as described previously^14^. As needed, 1 mM ferrous iron and 2 mM NTA, 100 *μ*M ferrous iron, or 100 *μ*M ferric malate was added to the anaerobic growth medium.

*R. palustris* strains were grown at 30°C aerobically in the dark in YP medium (0.3% yeast extract and 0.3% peptone) or anaerobically in photoheterotrophic medium (PM) supplemented with 10 mM succinate (PMS-10), as described previously^31^. Anaerobic cultures were incubated in a growth chamber with constant light (100 *μ*E of photosynthetically active radiation). As needed, 1 mM ferrous iron was added to the anaerobic growth medium. Because *R. palustris* can oxidize ferrous iron, 3.4 mM citrate trisodium dihydrate was added to prevent ferric iron precipitates from accumulating in the growth medium.

*E. coli* strains were grown aerobically at 37°C in LB or anaerobically at 30°C in M9 minimal medium supplemented with 0.4% glucose and 10 mM fumarate. For anaerobic growth, 285 *μ*M L-cysteine was added as a reducing agent. As needed, the anaerobic medium was supplemented with iron (1 mM ferrous iron and 2 mM NTA) or without iron (0.1 mN HCl and 2 mM NTA.

Antibiotics and selective reagents used are as follows: kanamycin (50 *μ*g/mL for *E. coli* and *S. putrefaciens* strains, 125 *μ*g/ml for *D. magneticus*, and 200 *μ*g/ml for *R. palustris*), streptomycin (50 *μ*g/ml for *E. coli* and *D. magneticus* strains), diaminopilmelic acid (DAP) (300 *μ*M for *E. coli* WM3064), and sucrose (10% for *R. palustris* and *S. putrefaciens*, 1% for *D. magneticus*).

### Plasmids and cloning

Plasmids used in this study are listed in Supplementary Table 10. In-frame deletion vectors targeting *fez*_*Rp*_ and *fez*_*Sp*_ were constructed by amplifying upstream and downstream homology regions from *R. palustris* CGA009 and *S. putrefaciens* CN-32 genomic DNA, respectively, using the primers listed in Supplementary Table 11. The homology regions were then inserted into the SpeI site of pAK31 using the Gibson cloning method. The deletion vector for *fezBC*_*Dm*_ was constructed by amplifying upstream and downstream homology regions from *D. magneticus* AK80 genomic DNA using the primers listed in Supplementary Table 11. The *P*_*npt*_-*strAB* cassette was subsequently ligated between the upstream and downstream homology regions of the deletion vector via BamHI. Expression plasmids for *fez*_*Rp*_ and *fez*_*Sp*_ were constructed by amplifying the respective gene cluster using the primers listed in Supplementary Table 11. The amplified DNA was inserted into HindIII/SpeI-digested pAK22 via the Gibson cloning method. The Δ*fezBC*_*Dm*_ complementation vector was constructed by amplifying *P*_*fez*_-*fezABC* from *D. magneticus* genomic DNA using the primers listed in Supplementary Table 11. The amplified DNA was then ligated into the SalI/XbaI sites of the expression vector pBMK7.

Plasmids were transformed into *E. coli* WM3064 and then transferred to *D. magneticus*, *S. putrefaciens*, or *R. palustris* via conjugation. For *D. magneticus*, conjugations were performed as described previously^9^. Allelic replacement of *fezBC*_*Dm*_ (*dmr_28330-40*) with *strAB* was achieved with streptomycin selection and sucrose counterselection as described previously^8^. Attempts to delete *fezABC*_*Dm*_ were unsuccessful. For conjugal transfer of plasmids to *R. palustris*, strains were streaked onto 1.5% YP agar plates and incubated aerobically at 30°C for 5 days. Two to three days prior to conjugation, single colonies were inoculated into YP medium and incubated aerobically at 30°C, until an OD_660_ of 0.2-0.7. Mid-log cultures of *E. coli* WM3064 carrying the plasmid to be transferred were mixed with *R. palustris* and spotted on 1.5% YP agar plates containing 0.3 mM DAP. After 2-3 days of incubation at 30°C, transconjugants were selected on 1.5% YP plates containing 200 *μ*g/ml kanamycin. For conjugal transfer of plasmids to *S. putrefaciens*, overnight cultures of *E. coli* WM3064 carrying the plasmid to be transferred and *S. putrefaciens* were mixed and spotted on 1.5% LB containing 0.3 mM DAP and incubated aerobically at 30°C for 1 day. Transconjugants were selected with 50 *μ*g/ml kanamycin. Δ*fez*_*Rp*_ and Δ*fez*_*Sp*_ candidates were selected on 10% sucrose plates, screened for kanamycin sensitivity, and deletions were confirmed by PCR.

### *S. putrefaciens* growth tests

For aerobic growth tests, *S. putrefaciens* WT and Δ*fez*_*Sp*_ strains were grown aerobically overnight and passaged 1:200 into LB supplemented with ethylenediaminetetraacetic acid (EDTA) at the specified concentrations. Cells were incubated at 30°C with continuous shaking and growth was monitored by measuring the A_600_ in a Sunrise microplate reader (Tecan).

For anaerobic growth tests, *S. putrefaciens* WT and Δ*fez*_*Sp*_ strains carrying a control plasmid and the Δ*fez*_*Sp*_ strain carrying the complementation plasmid (Δ*fez_Sp_∷fez_Sp_*) were inoculated in anaerobic LB supplemented with lacate, fumarate, 100 *μ*M ferrous iron, and kanamycin. Stationary phase cultures were then passaged 1:200 into anaerobic LB supplemented with lactate, fumarate, and kanamycin. EDTA and/or ferrous iron were supplemented at the specified concentrations. Cultures were incubated at 30°C and growth was monitored by measuring the A_600_ in a Sunrise microplate reader (Tecan) inside an anaerobic glovebag. Doubling times and lag times were calculated from the growth curves of raw values for individual strains shown in Extended Data Fig. 9b-e.

### Ferrosome isolation

*D. magneticus* was grown anaerobically in IL-RGM. Cells were then passaged 1:400 in two liters of anaerobic IL-RGM, as described above. When the culture reached an OD_650_ ~0.1, 100 *μ*M ferric malate was added. After three hours, cells were pelleted at 8,000xg for 20 minutes and flash froze in liquid nitrogen before storing at −80°C. Samples were observed by TEM before and after the addition of iron to ensure ferrosomes had formed. We found that this method enriches for both ferrosomes and magnetosomes (Extended Data Figure 2a-c). In order to prevent contamination with magnetosomes and magnetosome proteins, we isolated ferrosomes from a magnetosome gene island deletion strain, ΔMAI, and prepared the samples for proteomics.

Cell pellets were thawed on ice and resuspended with ice-cold LyA buffer (10 mM Tris HCl pH 8.0, 50 mM NaCl, and 1 mM EDTA) containing 250 mM sucrose, 1 *μ*g/ml leupeptin and pepstatin A and 1 mM PMSF. Cells were lysed by passing through a French pressure cell three times. The lysate was then passed through a 0.2 *μ*m filter to remove unlysed cells. The filtered cell lysate was gently layered over a 65% sucrose cushion and centrifuged at 35,000 rpm at 4°C for 2h. The resulting pellet was resuspended in 1 ml of LyA supplemented with leupeptin, pepstatin, and PMSF and washed two times with LyA before resuspending in a final volume of 50 *μ*l.

### Liquid Chromatography-Electrospray Ionization-Mass Spectrometry

Isolated ferrosomes (5 *μ*g) and whole cell lysate (50 *μ*g) were prepared for liquid chromatography-electrospray ionization-mass spectrometry (LC-ESI/MS). Each sample was combined with 0.06% RapiGest SF surfactant (Waters Corporation, Milford, MA) and 12 mM NH_4_CO_3_ pH 7.5 at 80°C for 15 minutes. Samples were incubated with 2.9 mM dithiothreitol at 60°C for 30 minutes followed by addition of 7.9 mM iodoacetamide (Sigma-Aldrich, St. Louis, MO) at room temperature for 30 minutes. Samples were then digested with 1:50 trypsin-protein (Promega) at 37°C overnight in the dark. Following digestion, 0.5% trifluoroacetic acid (Sequanal Grade, Thermo Fisher Scientific) was added and incubated at 37°C for 90 minutes to hydrolyze the RapiGest. The samples were centrifuged at 14,000 rpm at 4°C for 30 minutes and the supernatant was tranfered to a Waters Total Recovery vial (Waters Corporation, Milford, MA).

Trypsin-digested samples were analyzed at the QB3 Mass Spectrometry Facility using an Acquity M-class liquid chromatograph (LC) that was connected in-line with a Synapt G2-Si high-definition ion mobility mass spectrometer equipped with an electrospray ionization (ESI) source (Waters, Milford, MA). Mass spectrometry data analysis was performed using Progenesis QI for Proteomics software (Nonlinear Dynamics/Waters, Milford, MA) for relative protein quantification using a label-free approach.

### Electron microscopy

Whole-cell transmission electron microscopy was performed as described previously^6^. All TEM was done using the Tecnai 12 at the EM-Lab at the University of California, Berkeley.

### Multiple sequence alignments and tree construction

Unique protein sequences were obtained by searching DMR_28330, and selected subsequent target sequences, against all isolates in IMG/M ER^32^. Representative P_1B_-ATPase sequences from the characterized subgroups 1-4, CopA, ZntA, CopB, and PfeT, as well as a P_1A_-ATPase, KdpB, were also included. Sequences were aligned using MUSCLE in MEGA (7.0.26)^33^, with a gap open penalty of −6.9 and the resulting alignment was trimmed using Gblocks^34^. The trimmed alignment was used to generate a phylogeny using RAxML^35^ with the LG+G+F model (determined using SMS^36^) and 100 bootstraps. The tree was rooted with KdpB and visualized and annotated using iTol^37^.

To examine the synteny of *fez* gene clusters, we compiled a database of 304 FezB homologs identified in our phylogenetic analysis and the proteins encoded by the ten genes upstream and downstream of *fezB* for each species. We performed an all-versus-all search of these proteins using mmseqs2 10.6d92c^38^ (-s 7.5, -c 0.4, -e 1). The results from this search were uploaded into Cytoscape^39^ with an e-value cutoff <0.01 to generate a sequence similarity network. The Kyoto Encyclopedia of Genes and Genomes (KEGG)^40^ was used to identify conserved *fez* gene clusters containing FezB homologs (Extended Data Fig. 5a). These proteins were then mapped to nodes in eight different groups in the sequence similarity network. The Cytoscape plugin ClusterMaker^41^ was used to subdivide the following groups through Markov Clustering (MCL) with the inflation value set to 1.5. Group 1 (−log(e-value) 100); group 2 (−log(e-value) 2.5); and group 3 (−log(e-value) 5). Each group and subgroup with three or more proteins was then aligned with Clustal Omega^42^. For each alignment, HMMER 3.1b2 was used to build a hidden markov model which was searched against our database^43,44^. Subgroups that shared hits below a threshold of 1e^−20^ were merged and realigned. These alignments revealed a conserved GxxxG motif (or, less frequently, Gxxx[A/S] motif) for proteins in Groups 2 and 5 and an R-rich motif for proteins in Groups 1 and 3. Putative transmembrane domains were identified with TOPCONS 1.0^45^. Sequence logos of R-rich and GxxxG motif-containing proteins were generated with WebLogo^46^.

## Acknowledgements

We thank faculty at the EM-Lab at the University of California, Berkeley for their assistance with TEM; Anthony Iavarone at the QB3 Mass Spectrometry Facility for his help and advice; and Jeffrey Gralnick for *S. putrefaciens* CN-32. The QB3/Chemistry Mass Spectrometry Facility at the University of California, Berkeley receives support from the National Institutes of Health S10 Shared Instrumentation (grant number 1S10OD020062-01). Research reported in this publication was supported by funding from the National Institues of Health (R01GM084122 and R35GM127114), the Office of Naval Research (N000141310421), and the Bakar Fellows Program.

## Author contributions

C.R.G. and A.K. designed the study and wrote the manuscript. C.R.G. performed the experiments.

## Competing interests

The authors declare no competing interests.

## Additional information

**Extended data** is avalable for this paper.

**Supplementary information** is available for this paper.

## Extended Data

**Extended Data Fig. 1.**
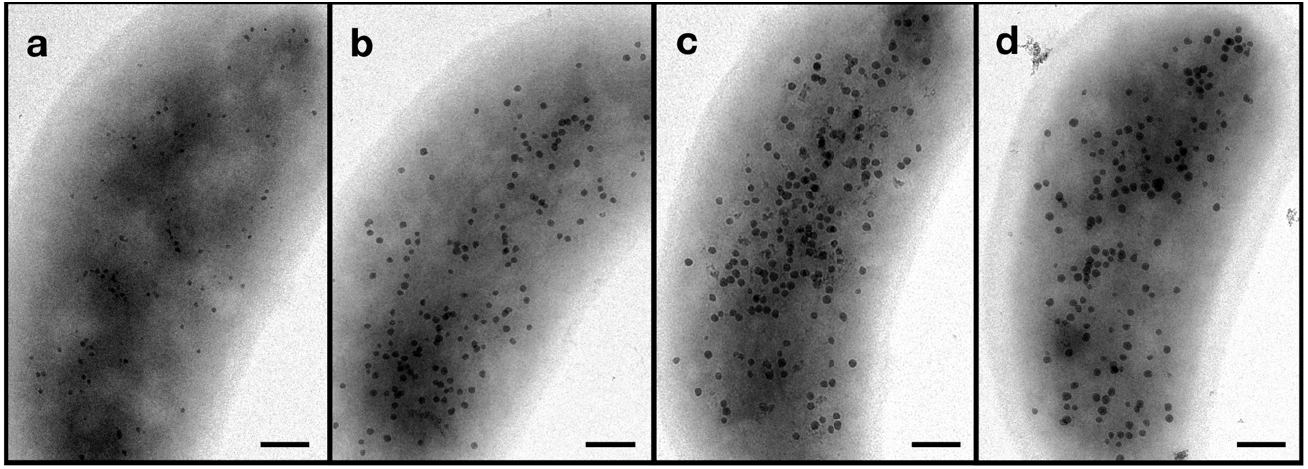
Ferrosomes are visible by TEM in whole *D. magneticus* cells upon recovery from iron starvation. In *D. magneticus*, ferrosomes are visible one hour after addition of 1 *μ*M (a), 10 *μ*M (b), 100 *μ*M (c), and 1 mM (d) ferric malate to iron-starved cells. Scale bars, 200 nm.

**Extended Data Fig. 2.**
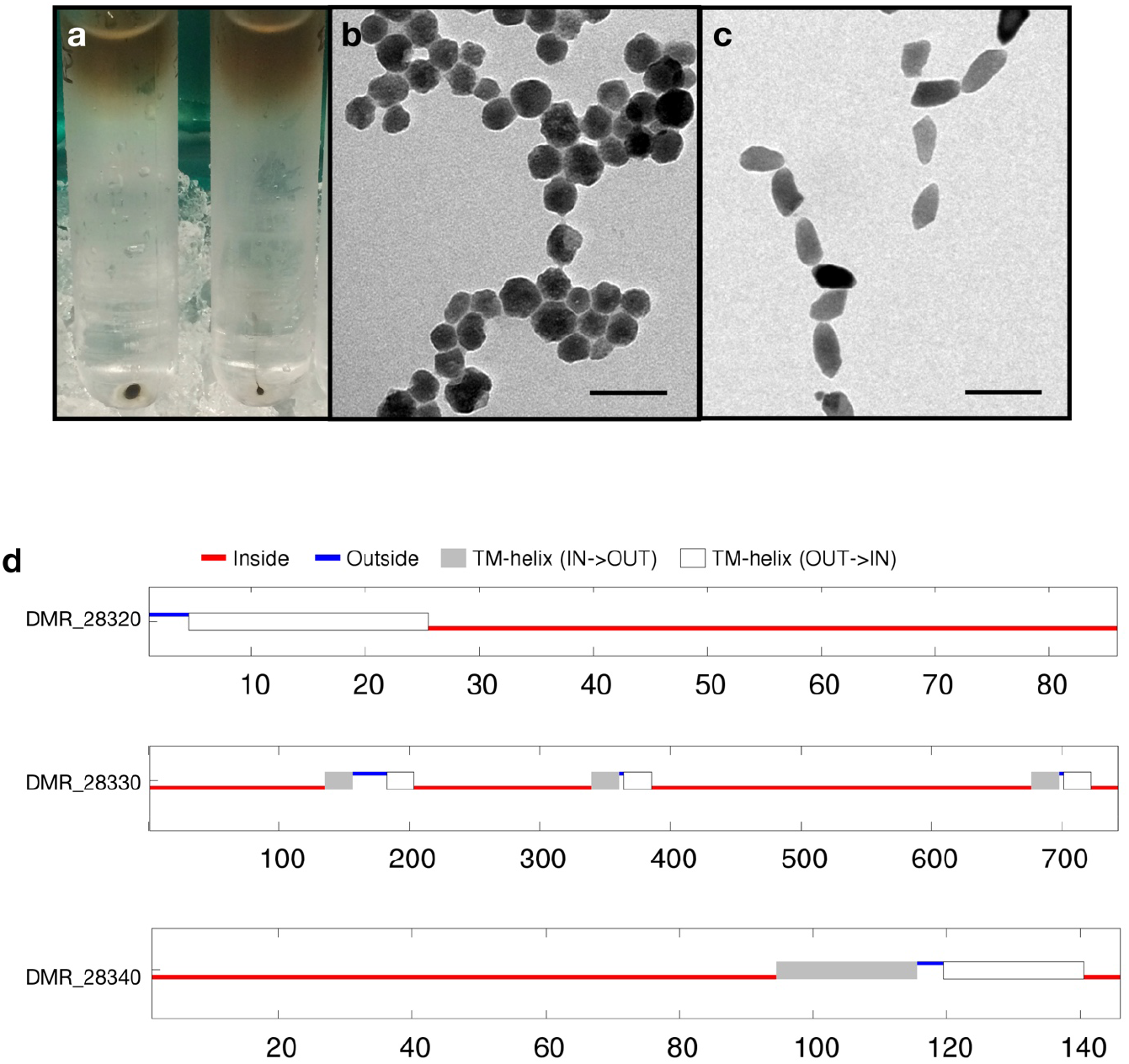
Isolation of ferrosomes. (a) Ferrosomes were isolated from ΔMAI *D. magneticus* cells that had transitioned from iron limited to iron replete medium by filtering whole cell lysate through a 65% sucrose cushion (left). A pellet is visible at the bottom of the sucrose cushion that contains ferrosomes, as confirmed by TEM (b). Magnetosomes were isolated from WT *D. magneticus* cells grown in iron replete medium using the same procedure (a, right; c). (b, c) Scale bars, 100 nm. (d) Membrane domain predictions of ferrosome-associated proteins in *D. magneticus*. DMR_28320 (FezA), DMR_28330 (FezB), and DMR_28340 (FezC) have 1, 6, and 2 putative transmembrane domains as predicted with TOPCONS 1.0.

**Extended Data Fig. 3.**
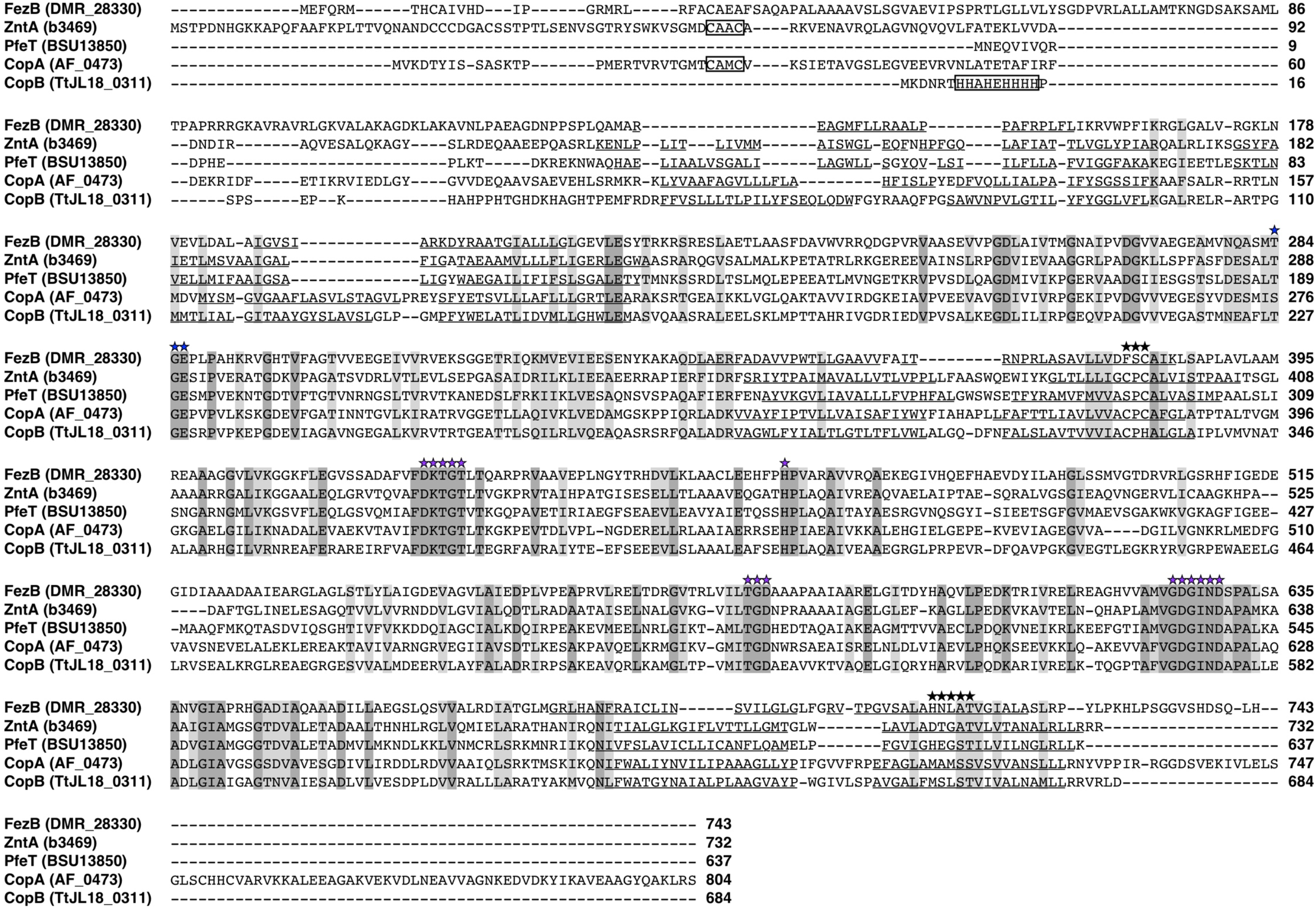
Multiple sequence alignment of FezB with characterized P_1B_-ATPases. Conserved functional motifs in the actuator domain and the ATP-binding domain are indicated with blue and purple stars, respectively. The CxxC and histidine-rich metal binding sites in the cytoplasmic N-terminal domain of ZntA, CopA, and CopB are boxed. Transmembrane regions, predicted using TOPCONS 1.0, are underlined for each sequence. Putative metal-binding sites in the transmembrane domains are indicated with black stars.

**Extended Data Fig. 4.**
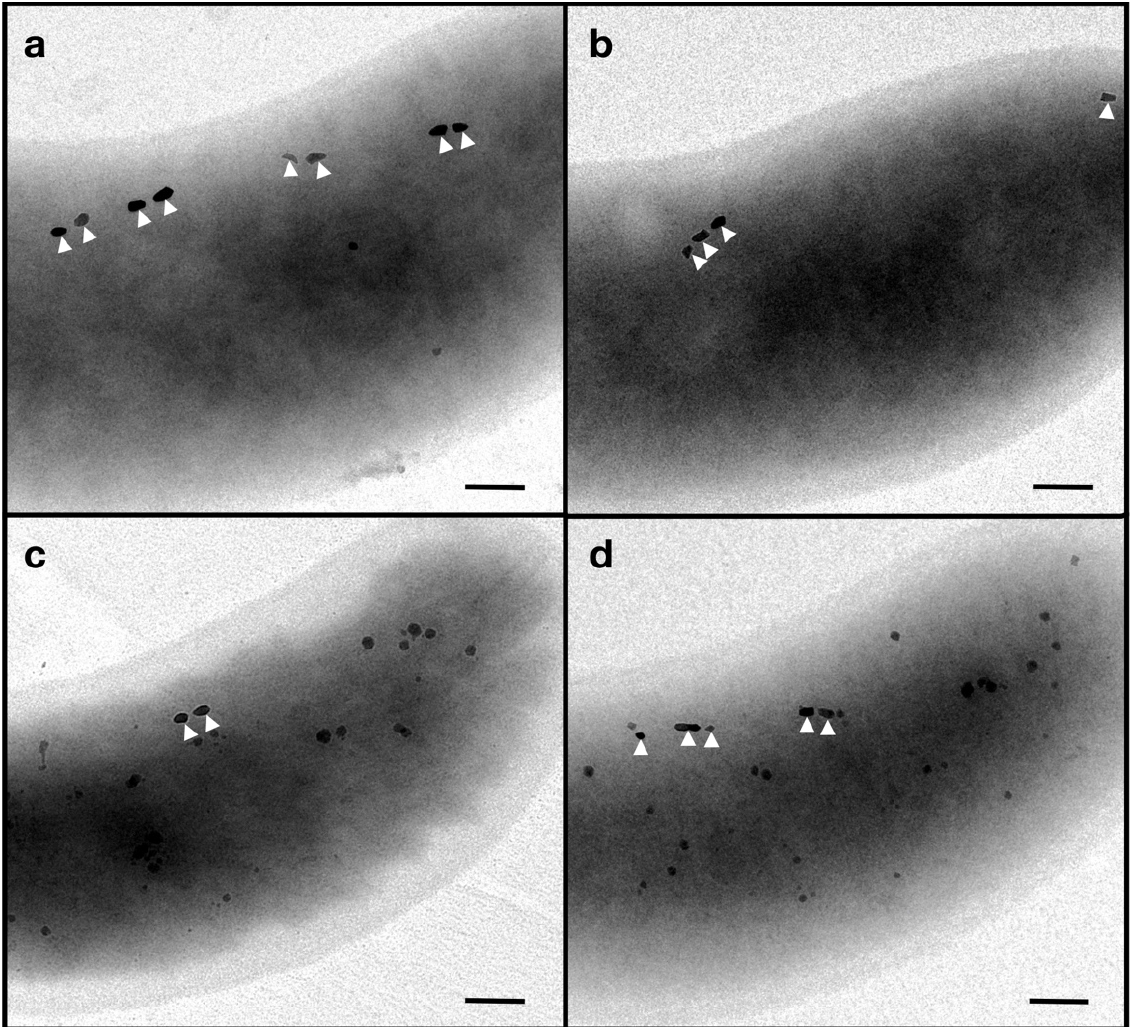
WT and Δ*fezBC_Dm_ D. magneticus* strains make ferrosomes in iron replete medium when expressing *fezAPC* in *trans*. Transmission electron micrographs of WT (a) and Δ*fezBC*_*Dm*_ (b) strains with a control plasmid make magnetosomes (white carets) when grown in iron replete medium. When expressing *fezABC* in *trans*, both the WT (c) and Δ*fezPC*_*Dm*_ (d) strains make magnetosomes as well as ferrosomes when grown in iron replete medium. Scale bars, 200 nm.

**Extended Data Fig. 5.**
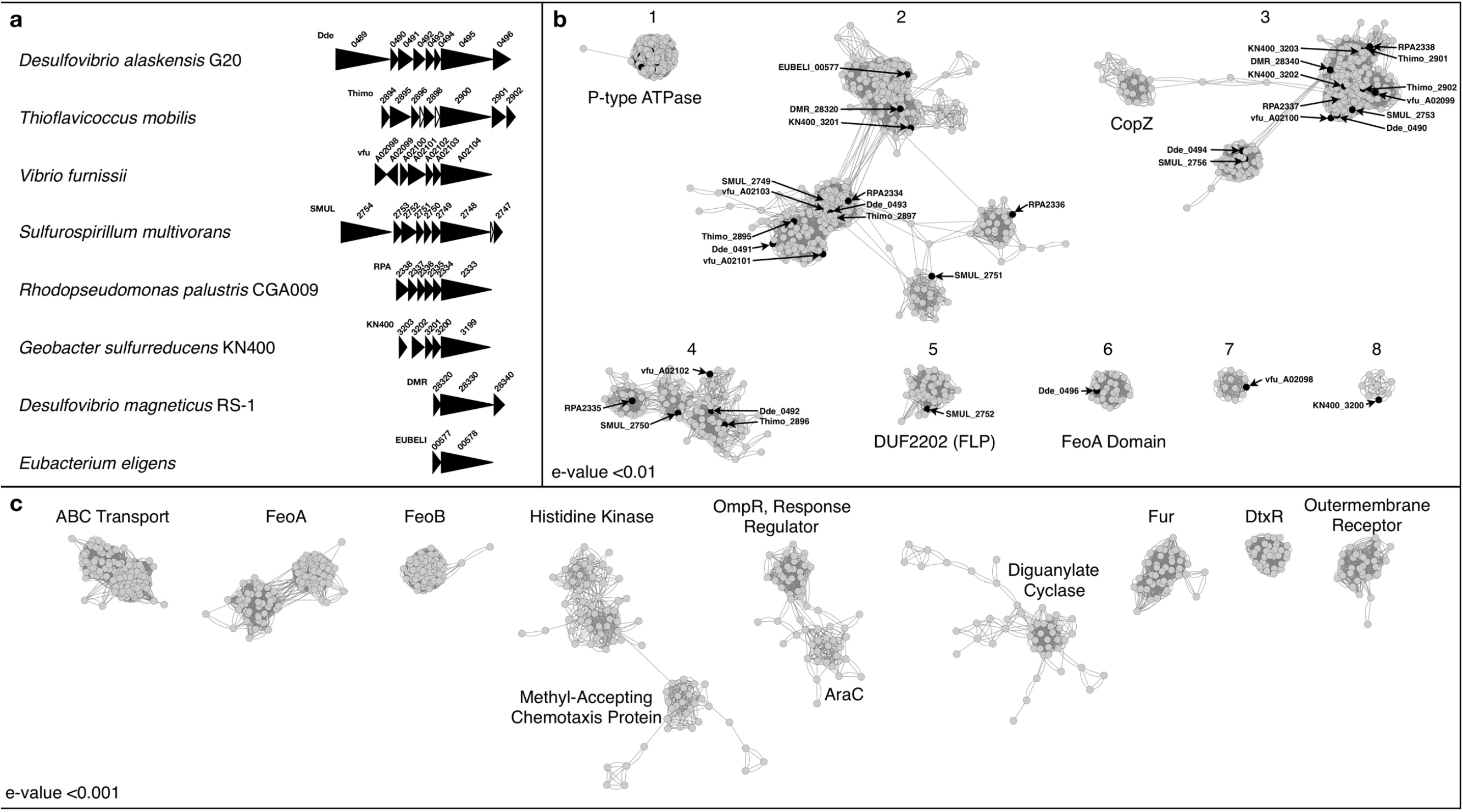
Sequence similarity network of proteins encoded by *fez* gene clusters and genes frequently found near *fez* gene clusters. (a) Conserved *fez* gene clusters that encode FezB homologs. Conserved genes within the clusters are colored black. Gene clusters were identified using the “Gene cluster” tool in KEGG for each FezB homolog (Dde_0495, Dde_0498, Thimo_2900, vfu_A02104, SMUL_2748, RPA2333, KN400_3199, DMR_28330, and EUBELI_00578). The second copy of FezB in *D. alaskensis*, Dde_0498, is not shown because it is not part of a conserved gene cluster. (b, c) Sequence similarity network highlighting the proteins encoded by ten genes upstream and downstream of 304 FezB homologs. Each node represents a protein and edges represent protein similarities that meet the specified e-value cutoff. (b) Network containing *fez* gene cluster-encoded proteins. Each group (labeled 1-8) contains one or more proteins encoded by conserved genes identified in (a) which are represented by black nodes and are labeled. Proteins or domains with an annotated function are labeled. Groups of proteins were further divided into subgroups which were used to identifiy proteins with GxxxG motifs in groups 2 and 5 and proteins with R-rich motifs in groups 1 and 3 (see Methods). The proteins represented in this network and their group/subgroup are listed in Supplementary Tables 3-6. (c) Network of proteins encoded by genes that are frequently found upstream and downstream of *fez* gene clusters. Only groups of more than 30 proteins are shown and the protein or domain annotation is labeled. Proteins with a known role in iron homeostasis are common and include iron transporters (FeoA, FeoB, outermembrane siderophore receptors, and some ABC transporters) and regulators (Fur and DtxR). The proteins represented in this network are listed in Supplementary Table 7.

**Extended Data Fig. 6.**
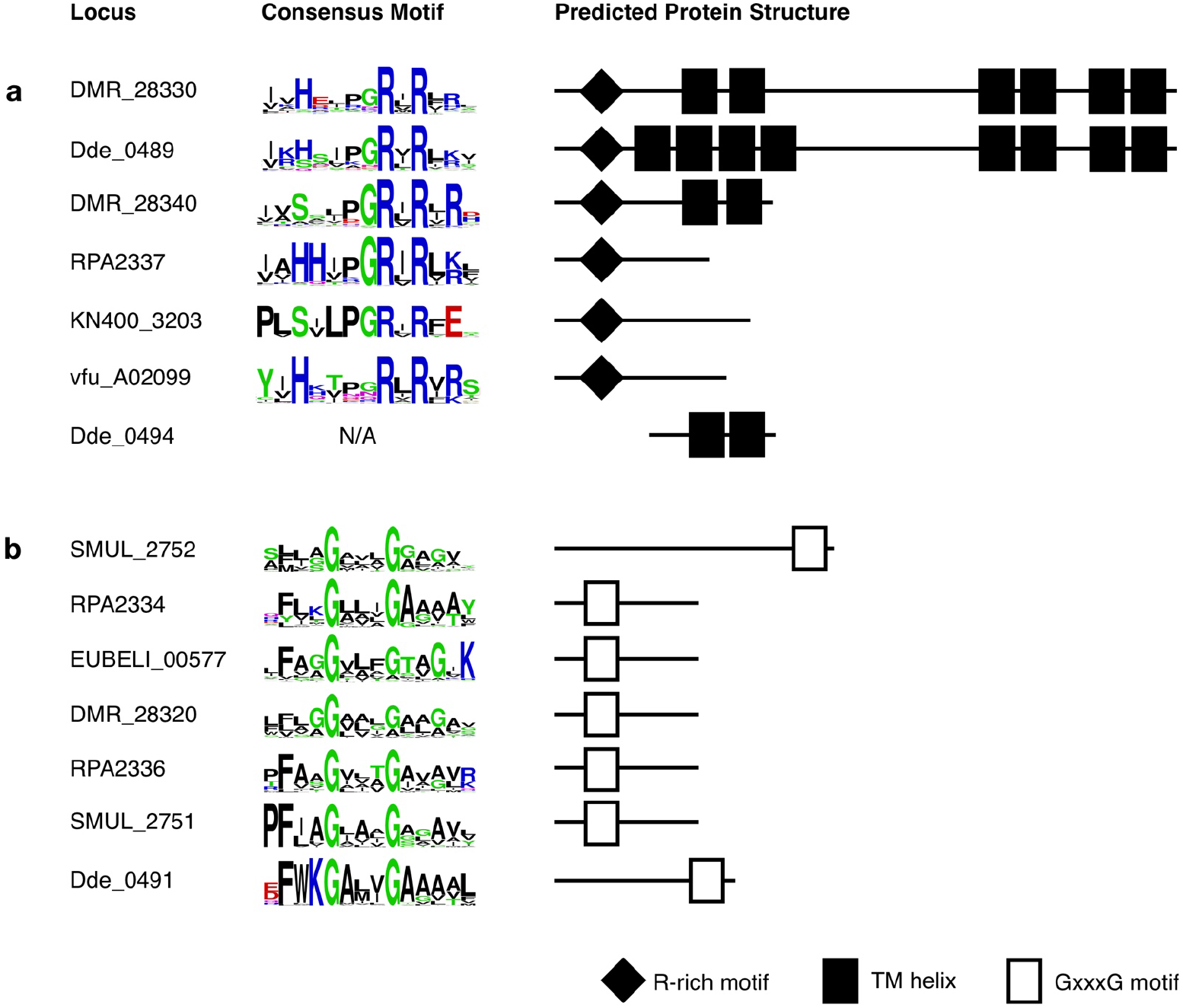
Representative proteins encoded by *fez* gene clusters with (a) an R-rich motif or (b) a GxxxG motif. Logo shows the consensus motif for the subgroup or group of proteins to which the representative protein belongs. Schematic shows the approximate location of the R-rich motif, putative transmembrane helices, and GxxxG motif for each protein (not to scale).

**Extended Data Fig. 7.**
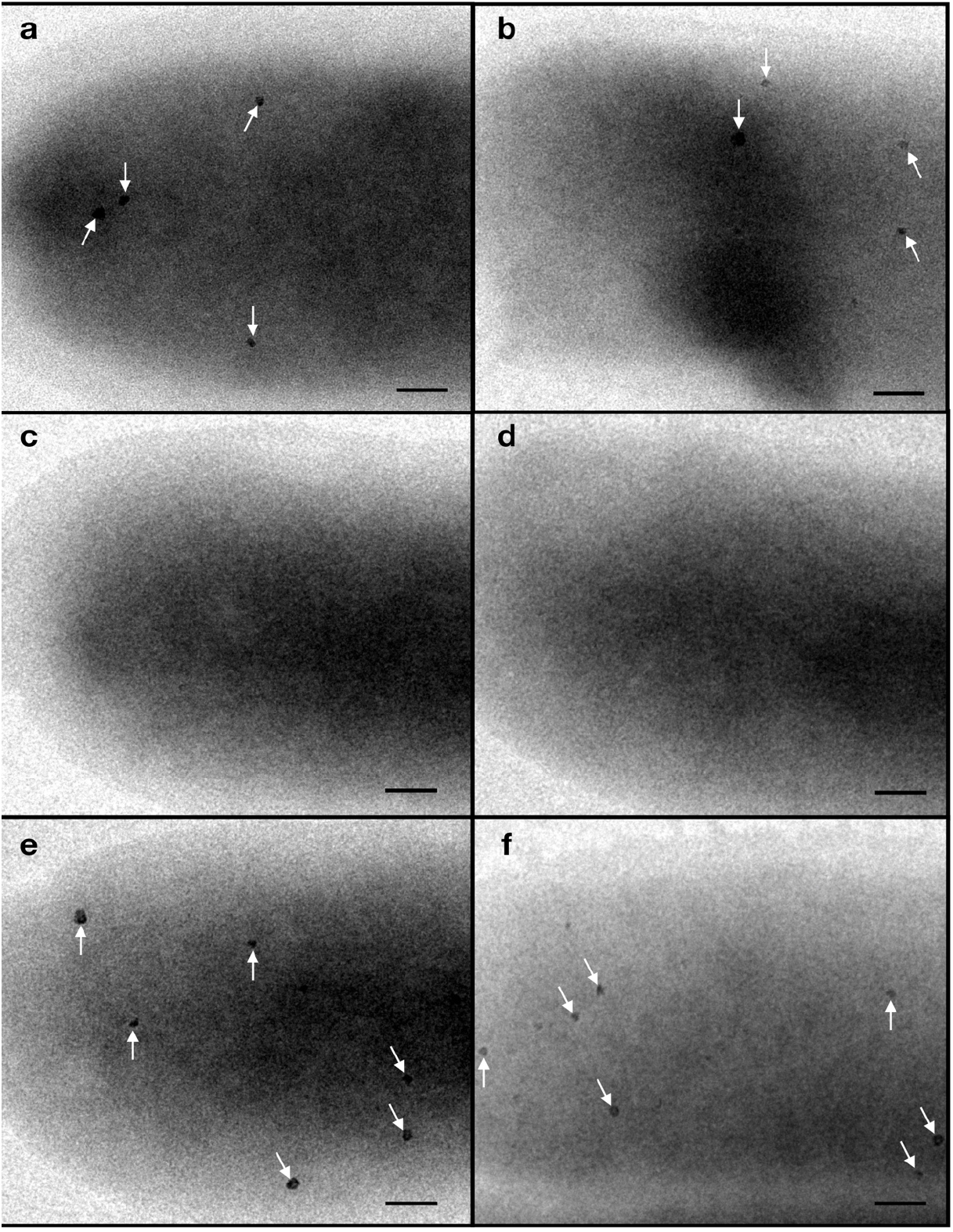
Transmission electron micrographs of *S. putrefaciens* strains respiring fumarate in medium supplemented with 100 *μ*M ferric malate (a, c, e) or 1 mM ferrous iron (b, d, f). Ferrosomes are found in WT *S. putrefaciens* (a, b), but not in the Δ*fez*_*Sp*_ mutant (c, d). This phenotype is rescued in the complementations strain Δ*fez_Sp_∷fez*_*Sp*_ (e, f). Scale bars, 100 nm.

**Extended Data Fig. 8.**
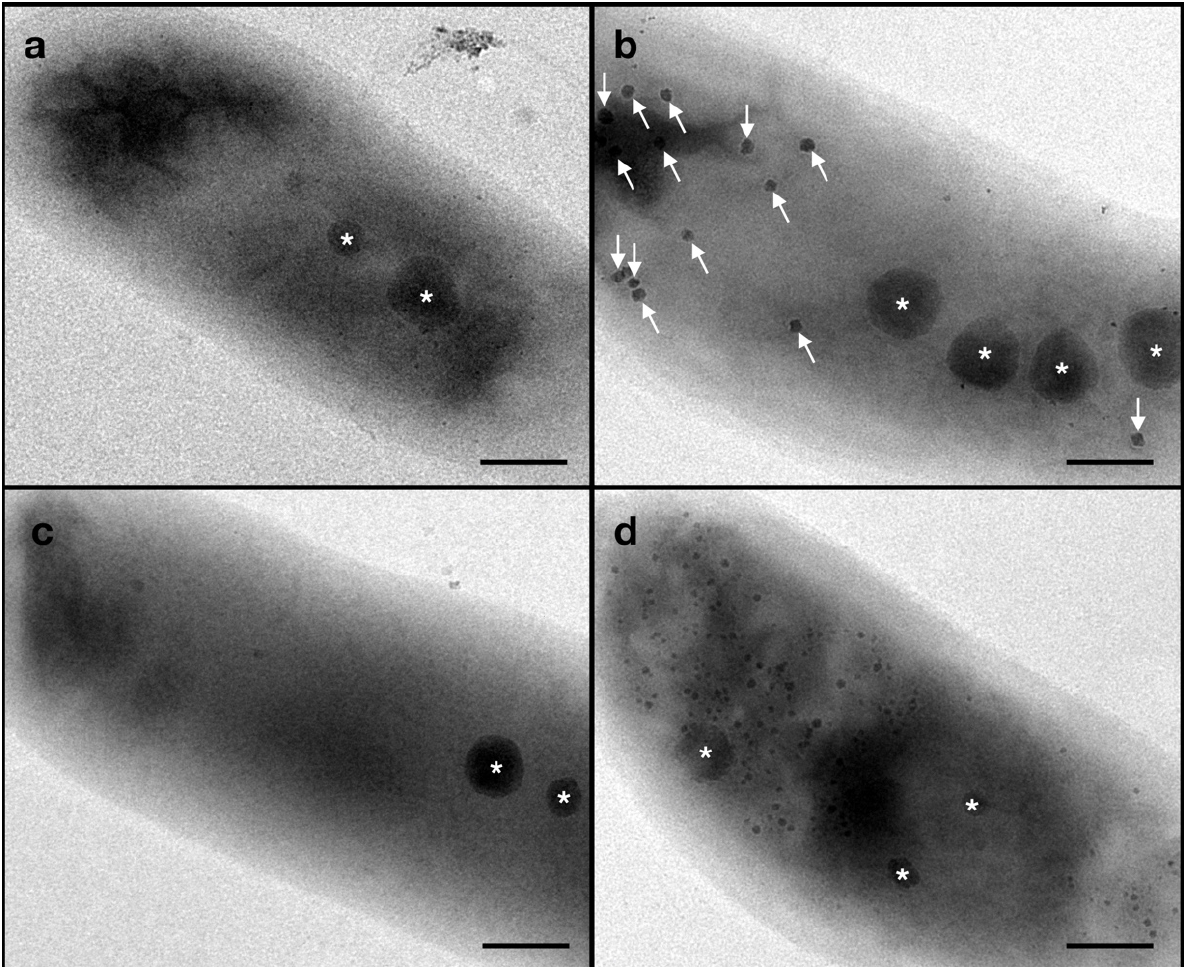
*fez* genes are essential for *R. palustris* to make ferrosomes. (a-d) Transmission electron micrographs of *R. palustris* CGA009. *R. palustris* CGA009 forms ferrosomes (white arrows) when grown anaerobically (b) and not aerobically (a). Deletion of the *fez*_*Rp*_ gene cluster abolishes ferrosome formation (c), a phenotype that can be complemented (d). Polyphosphate granules are indicated with a white asterisk. Scale bars, 200 nm.

**Extended Data Fig. 9.**
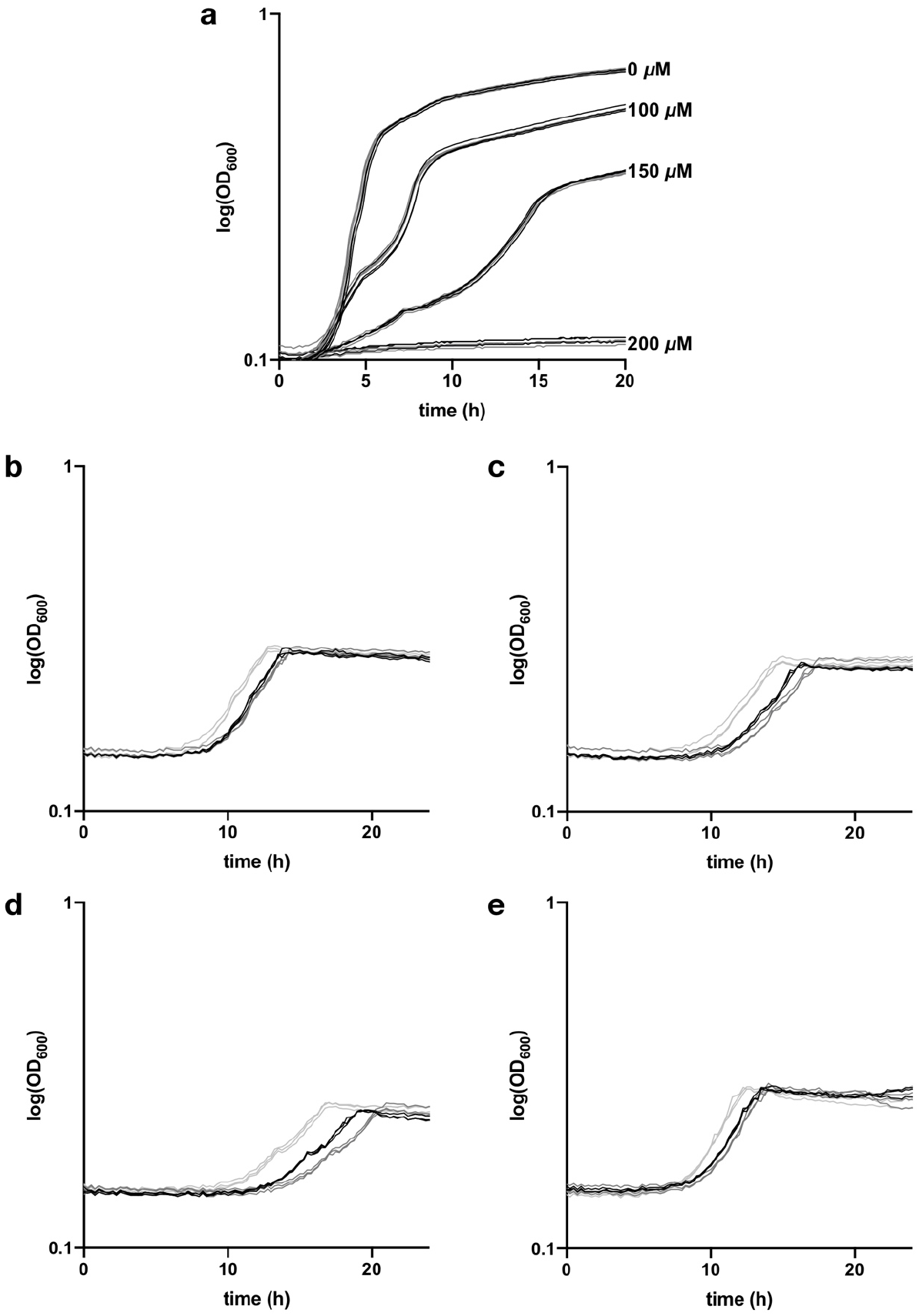
Effect of EDTA on the growth of *S. putrefaciens*. (a) Growth curves of the raw data for *S. putrefaciens* WT (black) and Δ*fez*_*Sp*_ (dark grey) grown aerobically with indicated concentration of EDTA. (b-f) Growth curves of the raw data for *S. putrefaciens* WT (black), Δ*fez*_*Sp*_ (dark grey) and Δ*fez_Sp_∷fez*_*Sp*_ (light grey) grown anaerobically with 0 *μ*M EDTA (b), 50 *μ*M EDTA (c), 100 *μ*M EDTA (d), and 100 *μ*M EDTA and 100 *μ*M ferrous iron (f).

## References

1. Andrews, S. C. Iron Storage in Bacteria. in Advances in Microbial Physiology (ed. Poole, R. K.) vol. 40 281–351 (Academic Press, 1998).

2. Touati, D. Iron and Oxidative Stress in Bacteria. Arch. Biochem. Biophys. 373, 1–6 (2000).

3. Andrews, S. C., Robinson, A. K. & Rodríguez-Quiñones, F. Bacterial iron homeostasis. FEMS Microbiol. Rev. 27, 215–237 (2003).

4. Andrews, S. C. The Ferritin-like superfamily: Evolution of the biological iron storeman from a rubrerythrin-like ancestor. Biochim. Biophys. Acta BBA - Gen. Subj. 1800, 691–705 (2010).

5. Nichols, R. J., Cassidy-Amstutz, C., Chaijarasphong, T. & Savage, D. F. Encapsulins: molecular biology of the shell. Crit. Rev. Biochem. Mol. Biol. 52, 583–594 (2017).

6. Byrne, M. E. et al. *Desulfovibrio magneticus* RS-1 contains an iron- and phosphorus-rich organelle distinct from its bullet-shaped magnetosomes. Proc. Natl. Acad. Sci. U. S. A. 107, 12263–12268 (2010).

7. Sakaguchi, T., Arakaki, A. & Matsunaga, T. *Desulfovibrio magneticus* sp. nov., a novel sulfate-reducing bacterium that produces intracellular single-domain-sized magnetite particles. Int. J. Syst. Evol. Microbiol. 52, 215–221 (2002).

8. Grant, C. R., Rahn-Lee, L., LeGault, K. N. & Komeili, A. Genome Editing Method for the Anaerobic Magnetotactic Bacterium *Desulfovibrio magneticus* RS-1. Appl Env. Microbiol 84, e01724–18 (2018).

9. Rahn-Lee, L. et al. A Genetic Strategy for Probing the Functional Diversity of Magnetosome Formation. PLOS Genet. 11, e1004811 (2015).

10. Argüello, J. M., Eren, E. & González-Guerrero, M. The structure and function of heavy metal transport P1B-ATPases. BioMetals 20, 233 (2007).

11. Smith, A. T., Smith, K. P. & Rosenzweig, A. C. Diversity of the metal-transporting P1B-type ATPases. J. Biol. Inorg. Chem. JBIC Publ. Soc. Biol. Inorg. Chem. 19, 947–960 (2014).

12. Padilla-Benavides, T., Long, J. E., Raimunda, D., Sassetti, C. M. & Argüello, J. M. A Novel P_1B_-type Mn^2+^-transporting ATPase Is Required for Secreted Protein Metallation in Mycobacteria. J. Biol. Chem. 288, 11334–11347 (2013).

13. Botella, H. et al. Mycobacterial P_1_-Type ATPases Mediate Resistance to Zinc Poisoning in Human Macrophages. Cell Host Microbe 10, 248–259 (2011).

14. Russ, W. P. & Engelman, D. M. The GxxxG motif: a framework for transmembrane helix-helix association. J. Mol. Biol. 296, 911–919 (2000).

15. Unterreitmeier, S. et al. Phenylalanine promotes interaction of transmembrane domains via GxxxG motifs. J. Mol. Biol. 374, 705–718 (2007).

16. Jarsch, I. K., Daste, F. & Gallop, J. L. Membrane curvature in cell biology: An integration of molecular mechanisms. J Cell Biol 214, 375–387 (2016).

17. Glasauer, S., Langley, S. & Beveridge, T. J. Intracellular Iron Minerals in a Dissimilatory Iron-Reducing Bacterium. Science 295, 117–119 (2002).

18. Glasauer, S. et al. Mixed-Valence Cytoplasmic Iron Granules Are Linked to Anaerobic Respiration. Appl. Environ. Microbiol. 73, 993–996 (2007).

19. VerBerkmoes, N. C. et al. Determination and Comparison of the Baseline Proteomes of the Versatile Microbe *Rhodopseudomonas palustris* under Its Major Metabolic States. J. Proteome Res. 5, 287–298 (2006).

20. Rey, F. E. & Harwood, C. S. FixK, a global regulator of microaerobic growth, controls photosynthesis in *Rhodopseudomonas palustris*. Mol. Microbiol. 75, 1007–1020 (2010).

21. Bose, A. & Newman, D. K. Regulation of the phototrophic iron oxidation (*pio*) genes in *Rhodopseudomonas palustris* TIE-1 is mediated by the global regulator, FixK. Mol. Microbiol. 79, 63–75 (2011).

22. Abdul-Tehrani, H. et al. Ferritin Mutants of Escherichia coli Are Iron Deficient and Growth Impaired, and fur Mutants are Iron Deficient. J. Bacteriol. 181, 1415–1428 (1999).

23. Rolfe, M. D. et al. Lag Phase Is a Distinct Growth Phase That Prepares Bacteria for Exponential Growth and Involves Transient Metal Accumulation. J. Bacteriol. 194, 686–701 (2012).

24. Bender, K. S. et al. Analysis of a Ferric Uptake Regulator (Fur) Mutant of *Desulfovibrio vulgaris* Hildenborough. Appl. Environ. Microbiol. 73, 5389–5400 (2007).

25. Uebe, R. et al. Deletion of a *fur*-Like Gene Affects Iron Homeostasis and Magnetosome Formation in *Magnetospirillum gryphiswaldense*. J. Bacteriol. 192, 4192–4204 (2010).

26. Wang, Q. et al. Physiological characteristics of *Magnetospirillum gryphiswaldense* MSR-1 that control cell growth under high-iron and low-oxygen conditions. Sci. Rep. 7, 2800 (2017).

27. Pereira, P. M. et al. Transcriptional response of *Desulfovibrio vulgaris* Hildenborough to oxidative stress mimicking environmental conditions. Arch. Microbiol. 189, 451–461 (2008).

28. Zhou, A. et al. Hydrogen peroxide-induced oxidative stress responses in *Desulfovibrio vulgaris* Hildenborough. Environ. Microbiol. 12, 2645–2657 (2010).

29. Caffrey, S. M. & Voordouw, G. Effect of sulfide on growth physiology and gene expression of *Desulfovibrio vulgaris* Hildenborough. Antonie Van Leeuwenhoek 97, 11–20 (2010).

30. Ho, T. D. & Ellermeier, C. D. Ferric Uptake Regulator Fur Control of Putative Iron Acquisition Systems in *Clostridium difficile*. J. Bacteriol. 197, 2930–2940 (2015).

## Methods References

31. Kim, M.-K. & Harwood, C. S. Regulation of benzoate-CoA ligase in *Rhodopseudomonas palustris*. FEMS Microbiol. Lett. 83, 199–203 (1991).

32. Chen, I.-M. A. et al. IMG/M v.5.0: an integrated data management and comparative analysis system for microbial genomes and microbiomes. Nucleic Acids Res. 47, D666–D677 (2018).

33. Kumar, S., Nei, M., Dudley, J. & Tamura, K. MEGA: A biologist-centric software for evolutionary analysis of DNA and protein sequences. Brief. Bioinform. 9, 299–306 (2008).

34. Castresana, J. Selection of conserved blocks from multiple alignments for their use in phylogenetic analysis. Mol. Biol. Evol. 17, 540–552 (2000).

35. Stamatakis, A. RAxML version 8: a tool for phylogenetic analysis and post-analysis of large phylogenies. Bioinformatics 30, 1312–1313 (2014).

36. Lefort, V., Longueville, J.-E. & Gascuel, O. SMS: Smart Model Selection in PhyML. Mol. Biol. Evol. 34, 2422–2424 (2017).

37. Letunic, I. & Bork, P. Interactive Tree Of Life (iTOL) v4: recent updates and new developments. Nucleic Acids Res. doi:10.1093/nar/gkz239.

38. Steinegger, M. & Söding, J. MMseqs2 enables sensitive protein sequence searching for the analysis of massive data sets. Nat. Biotechnol. 35, 1026–1028 (2017).

39. Markiel, S. P. et al. Cytoscape: a software environment for integrated models of biomolecular interaction networks. Genome Res. 13, 2498–504 (2003).

40. Kanehisa, M. & Goto, S. Kyoto Encyclopedia of Genes and Genomes. Nucleic Acids Res. 28, 27–30 (2000).

41. Morris, J. H. et al. *clusterMaker*: a multi-algorithm clustering plugin for Cytoscape. BMC Bioinformatics 12, (2011).

42. Sievers, F. et al. Fast, scalable generation of high-quality protein multiple sequence alignments using Clustal Omega. Mol Syst Biol 7, (2011).

43. Finn, R. D., Clements, J. & Eddy, S. R. HMMER web server: interactive sequence similarity searching. Nucleic Acids Res. 39, W29–W37 (2011).

44. Eddy, S. R. Accelerated Profile HMM Searches. PLOS Comput. Biol. 7, e1002195 (2011).

45. Bernsel, A., Viklund, H., Hennerdal, A. & Elofsson, A. TOPCONS: consensus prediction of membrane protein topology. Nucleic Acids Res. 37, W465–468 (2009).

46. Crooks, G. E. WebLogo: A Sequence Logo Generator. Genome Res. 14, 1188–1190 (2004).

